# Selective killing of plasma cell clones using splice-switching oligonucleotides targeting immunoglobulin variable exons

**DOI:** 10.1101/2024.08.30.610519

**Authors:** Jean-Marie Lambert, Antoine Praité, Marion Contini, Mohamad Omar Ashi, Anne Marchalot, Soazig Le Pennec, Christophe Sirac, Sandy Al Hayek, Laurent Delpy

## Abstract

Deregulated proliferation of a plasma cell (PC) clone is accompanied by an excess production of a monoclonal immunoglobulin (mo-Ig) component with unique V(D)J rearrangement features. In systemic Ig light chain (AL) amyloidosis, organ dysfunction is due to the deposition of amyloid fibrils composed of mo-Ig light chains in target tissues. Recent advances in exon skipping therapy using splice-switching antisense oligonucleotides (ASO) prompted us to examine a new antisense strategy targeting the variable (V) exon in myeloma cells. Indeed, we previously observed that the production of truncated Ig light chains, encoded by alternatively spliced mRNAs lacking V exon, heightened endoplasmic reticulum (ER) stress and triggered apoptosis of antibody-secreting plasma cells. We designed ASO hybridizing donor or acceptor V exon splice sites on mo-Ig pre-mRNAs. These compounds were very potent alternative splicing inducers and increased the production of V-domain-less truncated Ig chains. Remarkably, *in vitro* experiments and tumor xenograft models revealed that myeloma cells were highly sensitive to specific ASO treatment, compared to an irrelevant control-ASO. RNA-seq experiments further confirmed that the production of truncated Ig induced upon ASO treatment provoked a massive myeloma cell death through ER stress-associated apoptosis. In addition, high throughput sequencing of Ig repertoire demonstrated that ASO targeting JH2 or Jκ1 donor splice site induced massive elimination of VDJH2- or VJκ1-rearranged clones respectively, while sparing the others. Collectively, these data provide evidence that ASO targeting V exon on mo-Ig pre-mRNAs can emerge as new weapons to induce selective killing of plasma cell clones.

## INTRODUCTION

Plasma cells (PCs) are immunoglobulin (Ig) production factories that can secrete as many as 10^3^ Ig per second (Eyer et al., 2017; Khodadadi et al., 2019). The survival of antibody-secreting cells depends on their ability to alleviate endoplasmic reticulum (ER) stress associated with massive Ig synthesis. To ensure high Ig secretion, the unfolded protein response (UPR) is activated during PC differentiation. The UPR pathway adjusts ER capacity and promotes protein folding while increasing the proteasomal degradation of unfolded/misfolded proteins (Shaffer et al., 2004; Todd et al., 2009; Zhu et al., 2019). The three branches of the UPR are mediated by inositol-requiring protein-1 (IRE1), activating transcription factor-6 (ATF6) and protein kinase RNA-like ER kinase (PERK) (Ron and Walter, 2007). The PERK signaling pathway controlling translation through phosphorylation of the early initiation factor 2alpha (eIF2α) is suppressed during PC differentiation, whereas activation of the IRE1/ X-box binding protein 1 (XBP1) branch is critical (Gass et al., 2008; Reimold et al., 2001). IRE1α ensures the excision of 26-nt from *XBP1* mRNA, generating a short XBP1 isoform (XBP1s: XBP1 “spliced”) that regulates genes involved in protein synthesis, secretion and the ER-associated degradation (ERAD) pathway (Acosta-Alvear et al., 2007; Shaffer et al., 2004). Given that PC survival involves a tight regulation of intracellular protein homeostasis (proteostasis), modulating the ubiquitin-proteasome degradation with proteasome inhibitors (PI) stands as a gold standard therapeutic approach in patients with PC disorders (Auner and Cenci, 2015; Cohen et al., 2015; Gandolfi et al., 2017; Jaccard et al., 2014).

As a new checkpoint in PC homeostasis, disruption of proteostasis through alternative splicing of nonproductive Ig transcripts can generate truncated Ig polypeptides that impair PC differentiation by triggering ER stress-associated apoptosis (Srour et al., 2016). Indeed, the presence of a nonsense codon within the variable (V) exon activates a RNA surveillance pathway, referred as nonsense-associated altered splicing (NAS), which promotes skipping of the offending exon (Lambert et al., 2019, 2020). The extent of V exon skipping closely correlates with Ig gene transcription and is highly active in PC (Ashi et al., 2019). Consistent with the occurrence of a Truncated-Ig Exclusion (TIE-) checkpoint, mice expressing V-domain-less truncated Igκ light chains (ΔV-κLCs) exhibited a PC deficiency due to prolonged ER stress. Interestingly, the production of ΔV-κLCs affects both the expansion of PC after short-term immunization and the survival of long-lived PCs in bone marrow (Srour et al., 2016). Under physiological conditions, this production of ΔV-κLCs following NAS activation is limited to PC clones harboring biallelic Ig gene rearrangements and a nonsense V exon on the nonproductive allele.

Therefore, modulating Ig RNA splicing to produce truncated Ig polypeptides and selectively eliminate abnormal PC clones should stand as an attractive therapeutic approach for monoclonal gammopathies. Indeed, this strategy would be of particular interest in Ig light chain (AL) amyloidosis caused by the deposition of fibrillar aggregates composed of Ig light chain (LC) fragments in tissues, leading to organ dysfunction (Merlini et al., 2006). AL amyloidosis is one of the most common forms of systemic amyloidosis, mainly affecting the kidneys and heart, the latter of which is associated with a poor prognosis (Desport et al., 2012).

Antisense technology, particularly using splice-switching antisense oligonucleotides (ASOs) to modify RNA splicing has shown considerable promise, with significant results in treating type II spinal muscular atrophy (SMA) (Li, 2020; Hoy, 2017). Similar "gene surgery" has been developed in B-cell lineage to correct *Btk* (Bruton’s tyrosine kinase) splicing and restore its function in an animal model of XLA (X-linked gammaglobulinemia) (Bestas et al., 2014). Likewise, ASO-mediated skipping of exon 6 of *MDM4* (mouse double minute 4) reduces tumor growth in patient-derived xenograft (PDX) models of diffuse large B-cell lymphoma (DLBCL) (Dewaele et al., 2016). Since we recently demonstrated that morpholino oligos efficiently modulate Ig RNA splicing (Ashi et al., 2019; Marchalot et al., 2020; Mzarchalot et al., 2022), we examined the toxicity of ASO treatments targeting the V exon donor or acceptor splice sites (5’ss or 3’ss, respectively) in both normal and malignant antibody-secreting cells (ASCs). Given the crucial role of ER stress and protein homeostasis in PC survival, splice-switching ASOs targeting V exon on mo-Ig pre-mRNAs represents a promising therapeutic strategy to selectively induce apoptosis in malignant cells, potentially offering a new and effective treatment option for patients with AL amyloidosis and other monoclonal gammopathies.

## RESULTS

### ASO-mediated targeting of V exon 5’ss in malignant plasma cells

In monoclonal gammopathies, it has been suggested that the production of abnormal Ig impedes the proliferation of malignant PCs. According to this assumption, PCs from patients with AL amyloidosis exhibit cellular stress features, and transduced NS0 PC clones expressing amyloidogenic LCs showed slower growth rates compared to their nonamyloidogenic LC-expressing counterparts (Oliva et al., 2017). To investigate whether excessive synthesis of truncated Ig polypeptides is toxic to myeloma cells, we used antisense technology to modulate monoclonal Ig (mo-Ig) RNA splicing by targeting of V exon splice sites.

First, we specifically targeted the monoclonal *lambda* LC (*i.e. IGLV2-14*, and *IGLJ2*) expressed in RPMI 8226 cells with an ASO hybridizing the *IGLJ2* 5’ss on pre-mRNAs (ASO-Jλ2-5’ss) (**Fig.1A**). We observed a strong enhancement of alternative splicing in cells treated with ASO-Jλ2-5’ss, compared to an irrelevant control ASO (ASO-Ctrl). Quantification of full-length and alternative Igλ mRNAs showed that alternatively spliced transcripts (ΔV’-λLC) represented 44.7 ±24.4% and 62.9 ±21.2% of total mRNAs after 24 and 48h of ASO treatment, respectively (**Fig.1B**). Sequencing of ΔV’-λLC alternative transcripts identified a cryptic 5’ss within the complementary-determining region 1 (CDR1), encoding a truncated λLC polypeptide lacking the FR2 to FR4 region (**Fig.S1A**). According to Human Splicing Finder (HSF) splice site prediction algorithm (Desmet et al., 2009), this cryptic splice site is absent from the germline IGLV2-14 sequence, suggesting it was introduced by somatic hypermutation (SHM). Western Blot analysis showed that ΔV’-λLC mRNAs were translated into a truncated Ig of around 17 kDa (**Fig.1C**). Interestingly, treatment with ASO-Jλ2-5’ss for 48h was highly toxic to RPMI 8226 cells, resulting in approximatively 80% cell death compared to about 20% in control conditions (**Fig.1D**). Thus, ASO-mediated production of truncated ΔV’-λLC was correlated with increased cell death. Similar ASO-Jλ2-5’ss treatments were also performed on Igλ-expressing XG6 myeloma cells (IGLV1-40, and IGLJ2). Once again, we observed that ASO-mediated alternative splicing involved a cryptic 5’ss introduced by SHM, but located in the CDR2 (**Fig.1E and Fig.S1B**). Alternatively spliced ΔV’’-λLC mRNAs encoded a truncated Ig lacking the FR3 to FR4 region (**Fig.1F and Fig.S1B**). In contrast with data in **Fig.1D**, viability assays performed after ASO treatment indicated that the synthesis of ΔV’’-λLC polypeptides did not induce significant toxicity to XG6 cells **(Fig. 1G)**. One possible explanation could be that the production of truncated LCs with small V deletions could be tolerated by the cells. Therefore, the appearance of SHM-related cryptic 5’ss within the CDRs could strongly influence the pattern of ASO-mediated alternative splicing, precluding the targeting of V exon 5’ss in post-GC (germinal center) B cells.

**Figure 1:**
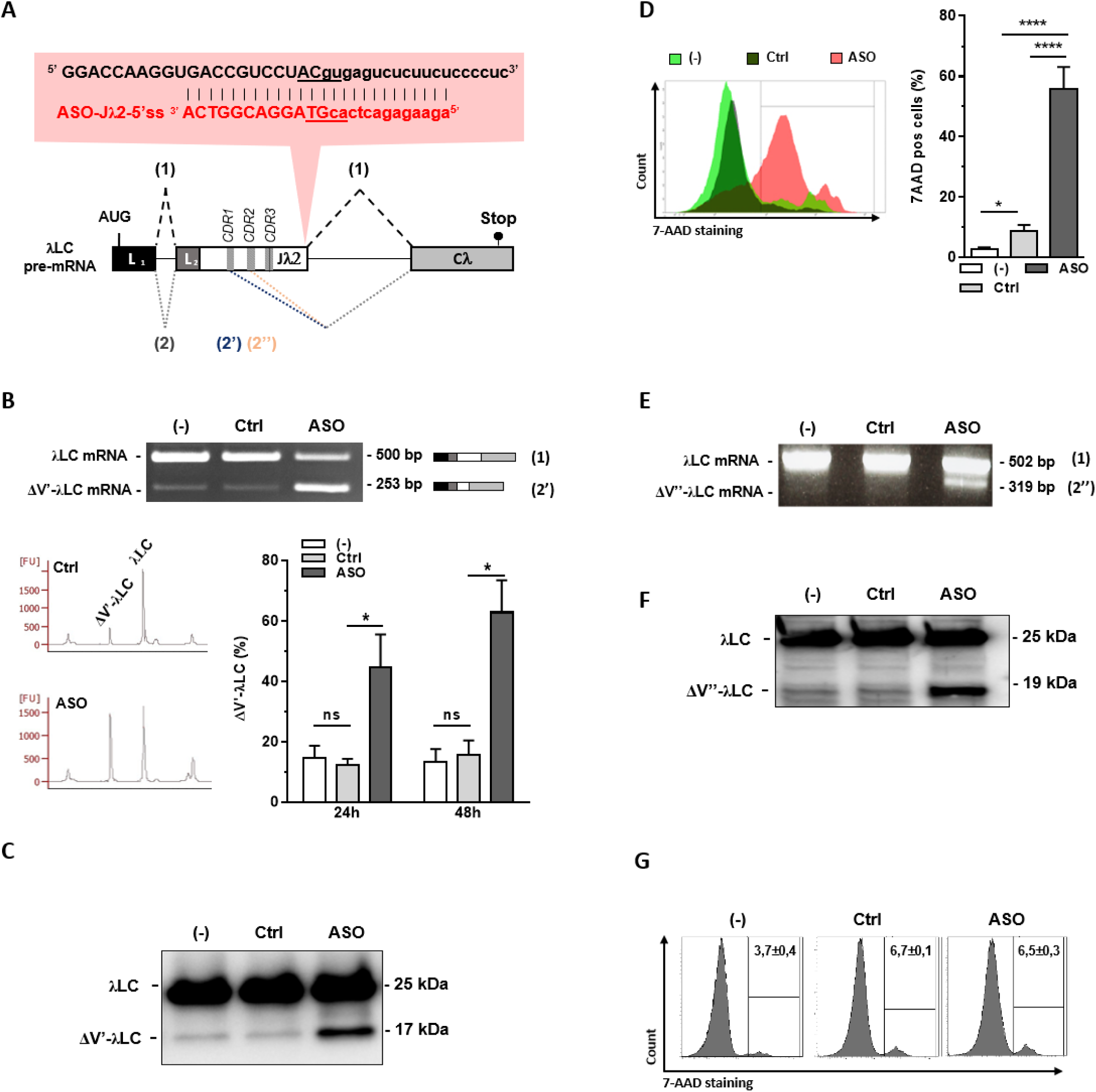
ASO targeting V exon 5’ss on monoclonal Igλ light chain pre-mRNAs. (A) RPMI 8226 cells expressed a monoclonal Igλ light chain with an *IGLV2-14* to *IGLJ2* rearrangement. The sequence of both *IGLJ2* 5’ss and specific ASO targeting this *IGLJ2* 5’ss (ASO-Jλ2-5’ss) are represented. ASOs were synthesized as "vivo-morpholino oligos" for passive administration (Gene Tools, LLC). (Uppercase: exon sequence; lowercase: intron sequence; L1: Leader part 1; L2: Leader part 2; Vλ: variable segment λ; Jλ2: joining segment λ2; Cλ: constant exon λ). Schematic representation of Igλ spliced transcripts are also depicted (dashed grey lines (1) represent normal splicing, and blue or orange (G) lines represent alternative splicing involving cryptic 5’ss (2)). **(B – G)** Experiments were performed in RPMI 8226 (**B-D**) and XG6 (**E-G**) cells treated or not with ASO (ASO: ASO-Jλ2-5’ss and Ctrl: ASO-Ctrl; 3 µM for 24 or 48 hours) **(B, top and E)** RT-PCR were performed using “Lpart1λ hs cons for” and “Cλ hs rev” primers to identify simultaneously identification of constitutive and alternative splicing of Igλ transcripts. PCR products were analyzed on agarose gels. Schematic representation of Igλ mRNAs is indicated on the right. **(B, bottom)** Relative quantification of amplification products was performed using Agilent 2100 Bioanalyzer. **(C and F)** Igλ light chain protein levels were analyzed by Western blot. **(D and G)** Percentage of 7-AAD-positive cells was determined by flow cytometry after 48h ASO treatment. Data from n=4-8 experiments are represented as mean ± SEM and unpaired two-tailed Student’s t test was used to determine statistical significance. (*P < 0.05,**P < 0.01, ***P < 0.001, ****P < 0.0001)

### Impact of ASO-mediated targeting of V exon 3’ss in myeloma cells

Consensus splice site profiling revealed that functional 3’ss generally have more constrained sequences compared to 5’ss. Consequently, ASO targeting regions around 3’ss generally resulted in more efficient exon skipping. To investigate this further, SK-MM-2 myeloma cells, which express a monoclonal *kappa* LC (*IGKV3-15*, and *IGKJ4*), were treated with an ASO targeting the 3’ss of V exon (ASO-Vκ3-15-3’ss)(**Fig.2A**). RT-PCR analysis and sequencing of alternative splicing transcripts revealed than ASO-Vκ3-15-3’ss strongly induced complete V exon skipping, compared to control conditions (**Fig. 2B**). Quantification of full-length and alternatively spliced *Igκ* mRNAs showed that ΔV-κLC mRNAs represented 67.7 ±7.1 % and 79.5 ±9.0 % of total Igκ mRNAs after 24 and 48h ASO treatment, respectively. At the protein level, ASO-Vκ3-15-3’ss treatment induced the production of an approximatively 12 kDa truncated ΔV-κLC (**Fig.2C**), while decreasing the amount of complete Igκ chains encoded by full-length κLC mRNAs (**Fig.S2**). Consistent with data in **Fig.1D**, the production of ΔV-κLC was correlated with increased cell death in SK-MM-2 cells treated with ASO-Vκ3-15-3’ss, compared to control conditions (**Fig. 2D**). Additionally, RNA-seq analysis of differentially expressed (DE) genes ((abs (log2FC) ≥ 0,75); pvalue Adj < 0,05) revealed 1100 upregulated and 524 downregulated genes in cells treated with ASO-Vκ3-15-3’ss compared to ASO-Ctrl (**Table S1 and Fig. 2E**). Interestingly, these differentially expressed genes were associated with pathways related to cell death, apoptosis, autophagy, protein ubiquitination and ER stress, suggesting that ER stress-associated apoptosis likely contribute to the massive cell death induced by ASO-Vκ3-15-3’ss treatment.

**Figure 2:**
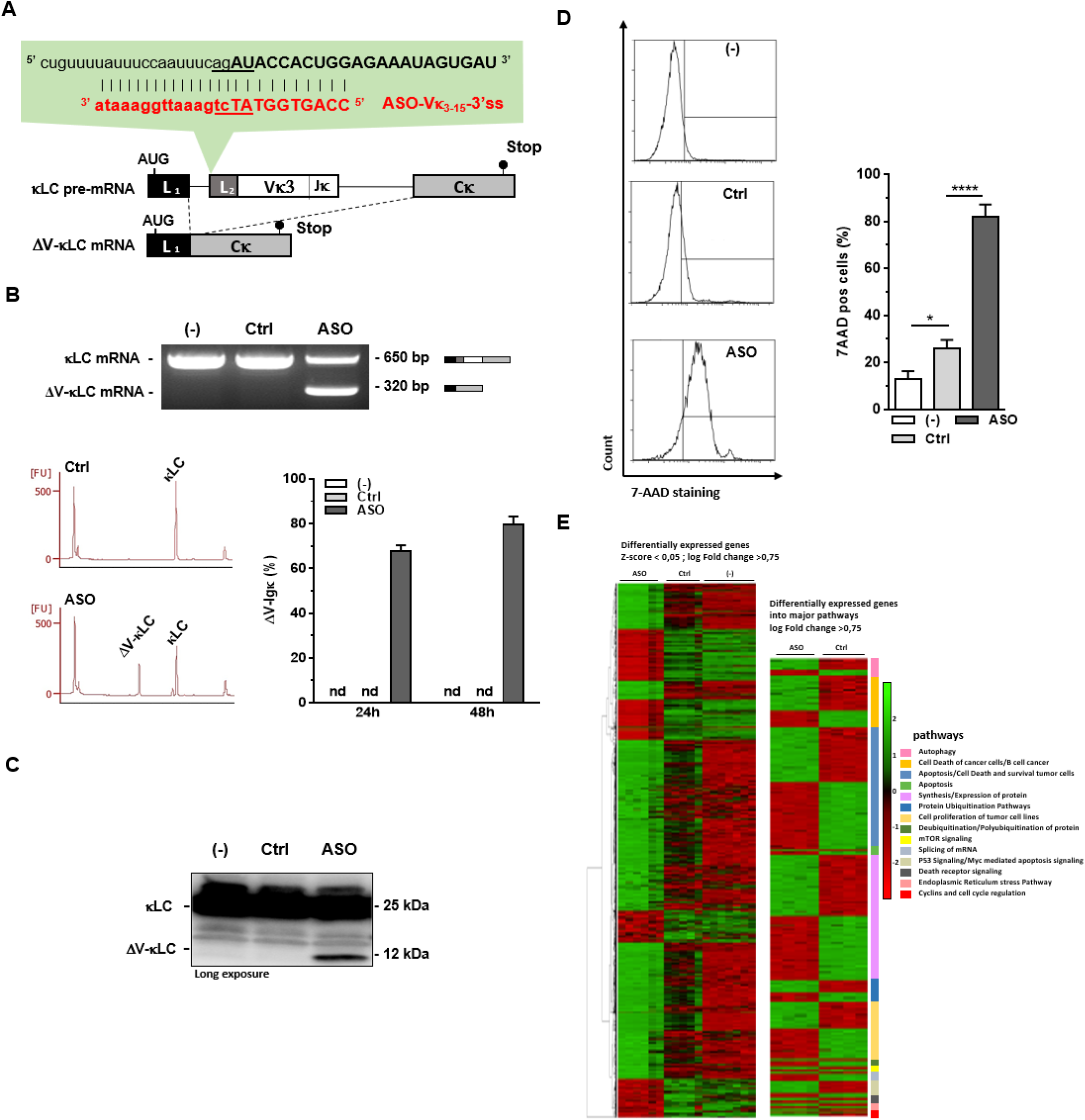
Exon skipping and production of harmful V-domain-less Igκ light chains using ASO targeting V exon 3’ss. **(A)** Sequence of "vivo-morpholino ASO" (ASO-Vκ3-15-3’ss) targeting *IGKV3-15* 3’ss on monoclonal Igκ pre-mRNAs expressed by SK-MM-2 myeloma cells (i.e. *IGKV3-15*, and *IGKJ4*). (Uppercase: exon sequence; lowercase: intron sequence; L1: Leader part 1; L2: Leader part 2; Vκ3: variable segment κ3; Jκ: junction segment κ; Cκ: constant exon κ). **(B – F)** Experiments were performed in SK-MM-2 cells treated or not with ASO (ASO: ASO-Vκ3-15-3’ss, Ctrl: ASO-Ctrl, (-): untreated; 3 µM for 24 or 48 hours) **(B, top)** RT-PCR were performed using “Lpart1 Vκ3 hs for” and “Cκ hs rev” primers to identify simultaneously constitutive and alternative splicing of Igκ transcripts. PCR products were analyzed on agarose gels. Schematic representation of Igκ mRNAs is indicated on the right. **(B, bottom)** Relative quantification of amplification products was done using Agilent 2100 Bioanalyzer. **(C)** Igκ light chain protein levels were analyzed by Western Blot. **(D)** Percentage of 7ADD-positive cells was determined by flow cytometry. **(E, left)** Heat map of RNA-seq analysis for the 1624 DE genes ((abs (log2FC) ≥ 0,75); p-value Adj < 0.05) from ASO, Ctrl and untreated groups. The differential analysis was performed using the following contrasts: Ctrl *vs* (-) and Ctrl *vs* ASO. The “DESeq2” package was used to perform the statistical modeling. **(E, right)** Among the differentially expressed genes identified in **Table S1,** the Ingenuity Pathway Analysis (IPA) software was used to represent as heat map specific genes involved in cell death and different proteostasis pathways. Data from n=4-7 experiments are represented as mean ± SEM and unpaired two-tailed Student’s t test was used to determine statistical significance. (*P < 0.05,**P <0.01, ***P < 0.001, ****P < 0.0001).

### ASO-mediated production of ΔV-κLC limits the growth of myeloma tumors *in vivo*

To evaluate the impact of ASO treatment on myeloma tumor growth *in vivo*, SK-MM-2 cells were engrafted subcutaneously into immunodeficient (Rag2^-/-^γC^-^) mice, and intratumoral (i.t.) injections of ASO-Vκ3-15-3’ss, ASO-Ctrl or PBS were administered once tumor volumes reached 100-200mm^3^ (**Fig. 3A up**). Remarkably, tumor growth was strongly reduced in mice that received 4 i.t. injections of ASO-Vκ3-15-3’ss (ASO), compared to those in control (ASO-Ctrl: Ctrl and PBS: (-)) groups (**Fig. 3A left**). Mice were ethically sacrificed when tumor volumes reached ∼800-900mm^3^ and animals treated with ASO-Vκ3-15-3’ss exhibited improved survival rates compared to the control groups (**Fig. 3A right**). Consistent with slower tumor growth combined to an altered production of complete Igκ, serum Igκ amounts were lower in mice treated with ASO-Vκ3-15-3’ss than in controls (**Fig.3B**).

**Figure 3:**
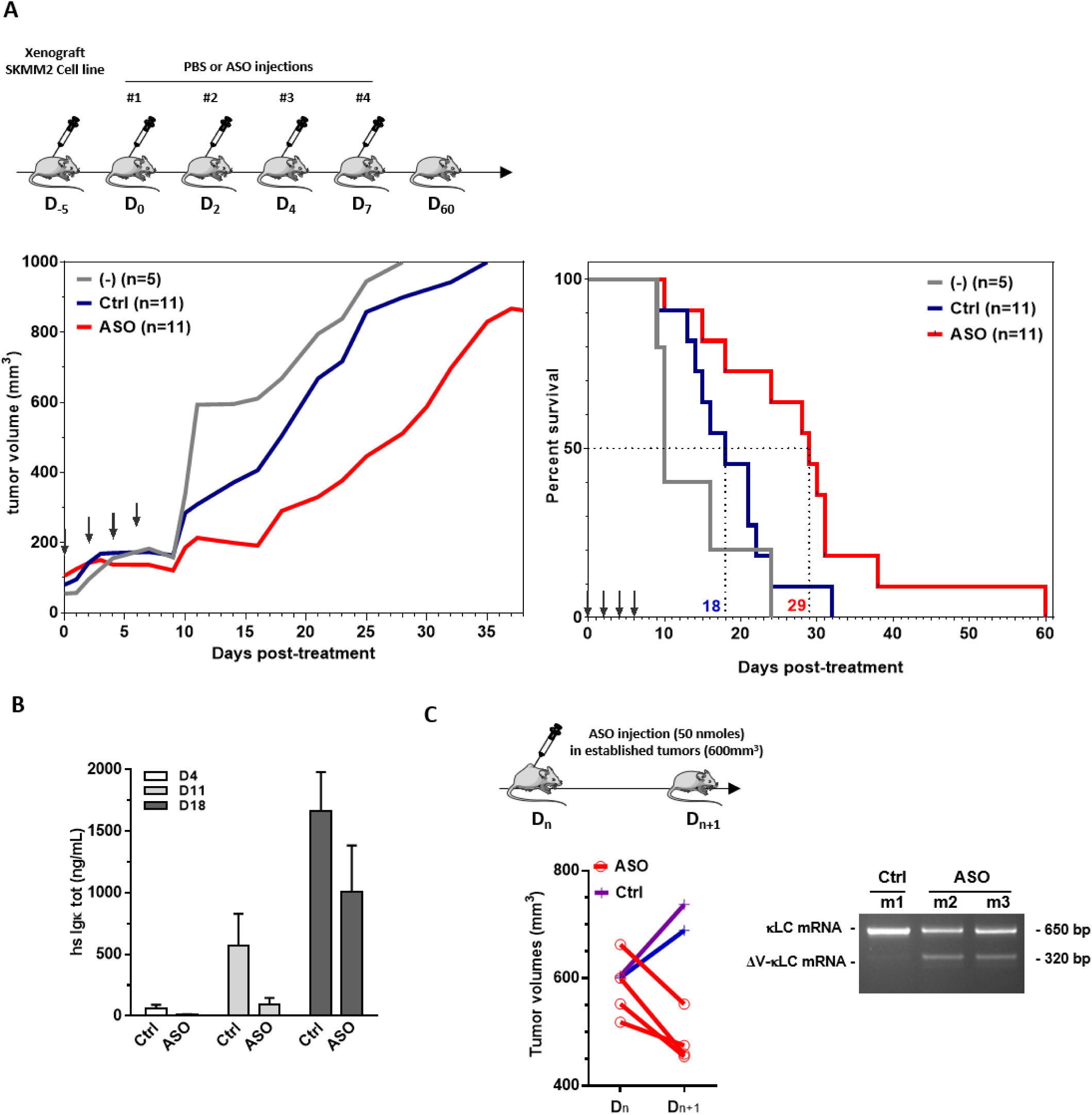
Antitumoral effect of ASO-inducing V exon skipping after engraftment of SK-MM-2 cells. **(A, top)** Representation of the experimental treatment protocol on mouse cohorts (ASO-Ctrl: Ctrl, n=11; ASO-Vκ3-15-3’ss: ASO, n=11; PBS, n=6). SK-MM-2 (2 x 10^6^ cells/mice) mixed with Matrigel were grafted in Rag2^-/-^γC^-/-^ by subcutaneous injection. The tumor size was calculated using the ellipsoid volume formula. When the tumor reached ∼100 mm^3^, mice received intertumoral injections of ASO (25 nmoles/injection) or PBS, 4 times every 2-3 days. The size of each tumour was measured daily and blood was collected every 4 days. **(A, bottom)** ASO injections are represented by arrows. The mean tumor volume (left) and the mouse survival curve (right) are shown (median survival: Ctrl 18 days, ASO 29 days, n = 11, * p = 0.03). **(B)** Concentrations of human Igκ in mouse mice sera (ELISA assays) at D4, D11 and D18. **(C)** A single injection of ASO (50 nmoles) was performed on established tumors (∼600mm^3^) and the tumor volume was determined at Dn and Dn+1. RNA was extracted from tumor cells after ASO treatment (Dn+1) and RT-PCR was performed using “Lpart1 Vκ3 hs for” and “Cκ hs”. Data from n=4-11 experiments are expressed as mean ± SEM and unpaired two-tailed Student’s t test was used to determine statistical significance. (*P < 0.05,**P < 0.01, ***P < 0.001, ****P <0.0001.)

We then aimed to evaluate the impact of ASO treatment on established tumors. A single i.t. injection of ASO was administered once the tumor volumes reached approximatively 600mm^3^. Tumor regression was observed in mice treated with ASO-Vκ3-15-3’ss for 24h, while tumors continued to grow after treatment with ASO-Ctrl (**Fig. 3C**). Exon skipping analysis of tumor cells injected with ASO further confirmed that tumor regression was associated with the presence of ΔV-κLC mRNAs (**Fig 3C**). These results demonstrate that ASO targeting of the V exon 5’ss effectively induces exon skipping and restricts myeloma tumor growth *in vivo*.

### Shaping of Ig repertoire using ASO targeting given V(D)J-rearranged RNA sequences

Using a dedicated mouse model harboring an extra-V exon in the *Igh* locus, we previously demonstrated that ASO treatment greatly induced V exon skipping in LPS-stimulated B cells, achieving over 80% efficacy in CD138-positive ASCs (Ashi et al., 2019). To investigate whether ASO-mediated V exon skipping and subsequent production of ΔV-κLC could selectively deplete ASCs, we stimulated splenic B cells with LPS in presence or absence of ASO targeting the germline mouse *IgkJ1* 5’ss (ASO-Jκ1-5’ss) (**Fig. 4A, left**). ASO-Jκ1-3’ss treatment resulted in complete V exon skipping in ASCs generated from naive B cells that had not undergone SHM (**Fig.4B**). Interestingly, rapid amplification of cDNA ends (RACE)-PCR analysis of Igκ repertoire revealed selective disappearance of full-length VκJκ1-rearranged transcripts in cells treated with ASO-Jκ1-3’ss, constituted only 2.6 ±1.5 % of total reads compared to 16.4 ±3.9% in cells treated with ASO-Ctrl (**Fig. 4C**). Similar results were obtained after analysis of VκJκ-rearranged clone frequency, with almost complete elimination of VκJκ1 clones (8.5 ±4.5%) compared to ASO-Ctrl (26.9 ±1.7 %) (**Fig. 4D**). These findings suggest that V exon skipping induced by ASOs targeting the J-segment 5’ss could shape the Ig repertoire, effectively mimicking the physiological TIE-checkpoint. Next, we performed similar experiments by using an ASO targeting *IghJ2* 5’ss (ASO-JH2-5’ss) (**Fig. 4A, right**). RACE-PCR Igµ repertoire analysis confirmed that ASO treatment blunted the differentiation of the vast majority of VDJH2-rearranged clones (**Fig. 4E and 4F**). Indeed, ASO-JH2-5’ss treatment of LPS-stimulated B cells resulted in a significant decrease in VDJH2 reads (6.75 ±4.0% of total reads), compared to control conditions (37.2 ±0.9 %) **(Fig. 4E)**, as well as decrease in the number VDJH2-rearranged clones (10.3 ±5.8%) relative to ASO-Ctrl (37.4 ±1.3 %) **(Fig. 4F)**. The selective elimination of ASC-expressing truncated V-less Ig heavy chain (HC) was consistent with previous findings for ER-stress associated apoptosis after production of ΔV-κLC (Srour et al., 2016). Therefore, it should be tempted to speculate that the TIE-checkpoint could broadly act on most PCs harboring biallelic V(D)J rearrangement of *Igh* and *Igκ* genes.

**Figure 4:**
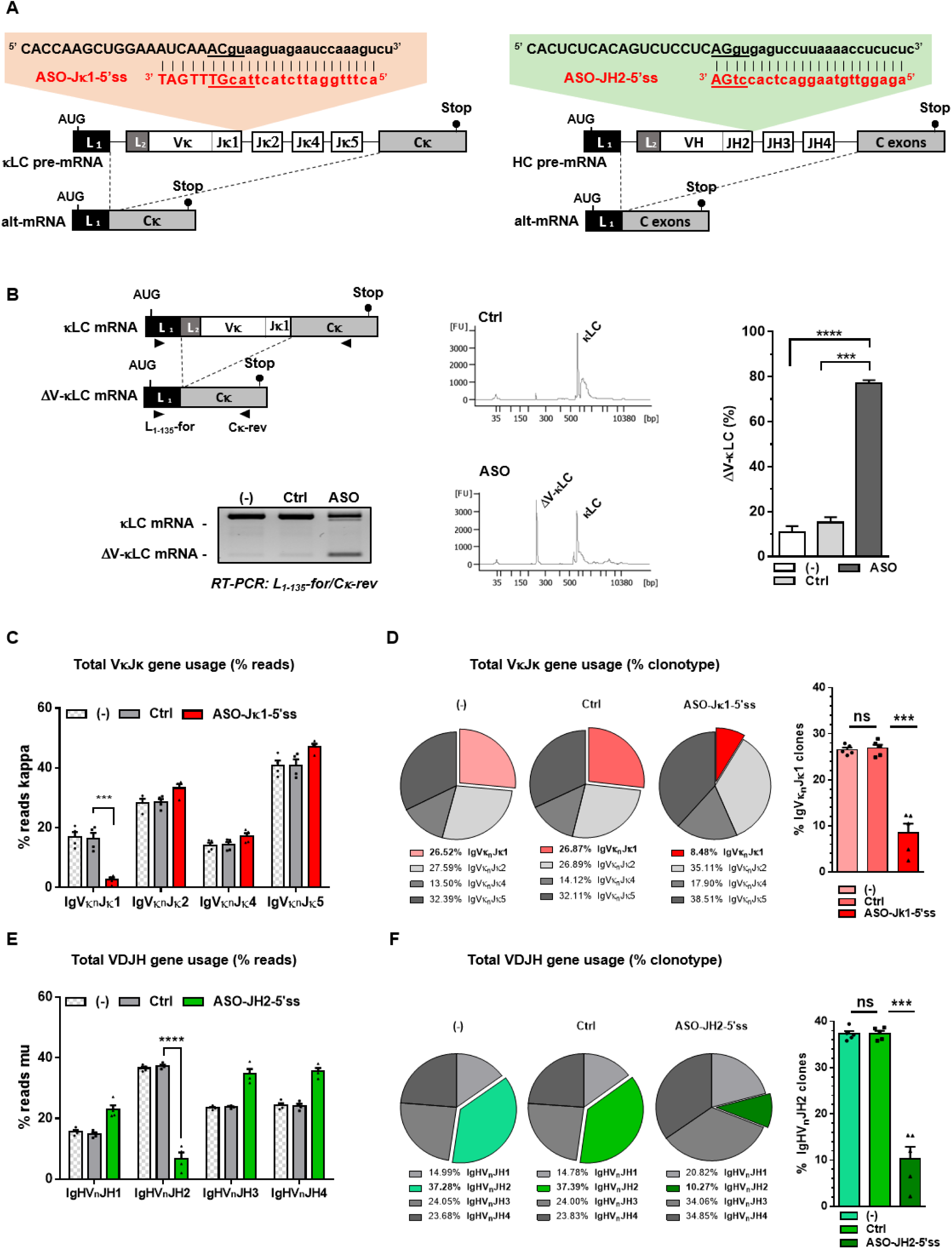
ASO Shaping of primary Ig repertoire using ASO targeting given J segment 5’ss. **(A, left)** Sequence of "vivo-morpholino ASO" (ASO-Jκ1-5’ss) targeting Jκ1 5’ss and, **(A, right)** sequence of ASO-JH2-5’ss targeting JH2 5’ss (uppercase: exon sequence; lowercase: intron sequence; L1: Leader part 1; L2: Leader part 2; Vκ/H: variable segment κ or H; Jκ/H: junction segment κ/H; Cκ/H: Constant exon κ/H). **(B – C)** Splenic B cells were isolated from C57BL/6 mice, stimulated with LPS for 4 days and treated with 2 μM ASO-Jκ1-5’ss, ASO-JH2-5’ss or an irrelevant control ASO (ASO-Ctrl), during the last 3 days **(B, left)** RT-PCR was performed using “Lpart1 Vκ1-135 mur for” and “Cκ mur rev” primers to identify constitutive and alternative splicing of Igκ transcripts simultaneously **(B, right)** Relative quantification of amplification products was done using the Agilent 2100 Bioanalyzer. **(C-F)** After RNA extraction, 5’RACE-PCR were performed using Cκ **(C-D)** or CH1γ **(E-F)** reverse primers and the clonal abundance was estimated after high-throughput sequencing of V(D)J junctions using the IMGT/HighV-QUEST software. Bar graphs represent the number of reads from VκJκ **(C)** and VDJH **(E)** junctions according to the presence of specific Jκ and JH segments, respectively **(D and F).** Pie charts and bar graphs represent the distribution of V(D)J clonotypes indicating a marked decrease in the frequency of VnJκ1 clones after treatment with ASO-Jκ1-5’ss **(D)** and VnDnJH2 clones after treatment with ASO-JH2-5’ss **(F)**. Data from n=4-6 experiments are expressed as mean ± SEM and unpaired two-tailed Student’s t test was used to determine statistical significance. (*P < 0.05,**P < 0.01, ***P < 0.001, ****P < 0.0001)

## DISCUSSION

In this study, we validated that V exon skipping can be effectively achieved by using ASO targeting V exon splice site on Ig pre-mRNAs. This opens new pathways for therapeutic activation of the TIE-checkpoint in monoclonal gammopathies (patent WO2017089359A1) (Srour et al., 2016). The production of truncated Ig chains induced by ASO treatment represents a pioneer therapeutic approach that should selectively eliminate tumor plasma cells. We demonstrated that ASO-mediated modulation of Ig pre-mRNA splicing leads to rapid cytotoxicity and subsequent apoptosis in myeloma cells.

Successful development of ASOs for specific targeting of monoclonal Ig expressed by tumor PCs relies on accurately identifying the V(D)J rearrangement and the intronic sequences flanking this V exon. Since tumor PCs are post-GC cells that exhibit extensive SHMs in their V regions and have typically undergone class switch recombination (CSR), precise sequencing of these regions is essential. Indeed, we have observed that mutations introduced by SHM can create or reinforce the predictive score of cryptic 5’ss within the CDRs, complicating the prediction of ASO-mediated alternative splicing pattern. However, targeting the V exon 3’ss harboring a more constrained consensus sequence due to the polypyrimidine tract, limits the use of unwanted cryptic splice sites, making it a highly effective strategy for inducing complete V exon skipping.

Our results obtained after engraftment of SK-MM-2 cells revealed slower tumor growth and improved median survival in the group treated with ASO hybridizing V exon 3’ss, compared to control conditions. The development of ASO compounds capable of targeting tumor PCs in medullary niches after systemic administration will be of great interest for the treatment of monoclonal gammopathies.

In AL amyloidosis, antisense strategies using siRNAs, or even phosphorothioate ASOs triggering RNase H1 cleavage of RNA targets, have been employed to target monoclonal Ig LCs (Ohno et al., 2002; Phipps et al., 2010; Hovey et al., 2011; Zhou et al., 2014; Ma et al., 2016). These strategies have been shown effectiveness in reducing monoclonal LC levels. In addition, research by Comenzo and colleagues demonstrated that pools of siRNAs targeting the *kappa* or *lambda* constant exon could induce the death of tumor PCs expressing complete Ig, composed of both HCs and LCs (Zhou et al., 2014; Ma et al., 2016). These studies showed that the reduction in LC levels following siRNA treatment caused protein stress, likely due to intracellular accumulation of unpaired HCs. Our strategy, which involves the production of truncated LCs lacking V domains that are intrinsically toxic to PCs offers a key advantage: it is applicable to all patients with AL amyloidosis, not just the subset (approximately 40 %) whose tumor PCs express complete (HC and LC) monoclonal Ig. Moreover, rather than the targeting of invariant constant exons, our approach targets the V exon specifically and is restricted to the monoclonal LC expressed by the tumor PCs. Therefore, our strategy could enable the selective elimination of tumor PC clones, while preserving the patient’s overall humoral immunity.

Improving the biodistribution of ASOs remains a major challenge across systemic antisense approaches beyond the liver. The development of novel ASO conjugates may allow efficient targeting of tumor PC niches in bone marrow. For example, antibody-ASO conjugates have already been developed in systemic diseases such as precursor B cell acute lymphoblastic leukemia (preB-ALL) (Satake et al., 2016). In monoclonal gammopathies, it would be interesting to couple ASOs targeting the V exon of monoclonal Ig to an anti-CD38 antibody, such as Daratumumab used in the clinic (van de Donk and Usmani, 2018). Other ASO conjugates have been explored and PPMO (peptide conjugated-phosphorodiamidate morpholino oligonucleotides) like B-PMOs (B peptide-conjugated PMO) could be useful to target B-lineage cells while ensuring a highly splice-switching efficiency (Bestas et al., 2014). We have recently shown that systemic injections of Pip6a-PMO conjugates (PMO internalization peptide 6a) are able to target ASCs *in vivo* (Marchalot et al., 2022).

Many questions remain regarding the impact of truncated Ig chains produced by alternative splicing in ASCs. Consistent with our previous findings revealing the occurrence of a TIE-checkpoint during normal PC differentiation (Srour et al., 2016), ASO-mediated production of V-domain-less truncated Ig should be considered as an interesting therapeutic approach for monoclonal gammopathies. Optimizing delivery methods to achieve precise biodistribution in bone marrow niches could accelerate the development of these innovative personalized therapies, enabling targeted elimination of tumor PC clones while sparing healthy cells.

## MATERIALS AND METHODS

### Cell culture

The human myeloma cell lines SK-MM-2 from DSMZ (Germany) and RPMI 8226 from ATTC (USA) were cultured in RPMI 1640 medium in the presence of 10% fetal calf serum (FCS). Patient-derived XG6 cells were kindly provided by J. Moreaux (IGH - UMR 9002 Montpellier, France) and in RPMI 1640 medium in the presence of 10% FCS and 5 ng/ml recombinant human IL-6. All cell lines were checked for the absence of mycoplasma (MycoAlert Mycoplasma Detection Kit; Lonza, Levallois, France). For ASO treatments, vivo-morpholino ASOs (ASO-Jλ2-5’ss: 5′-AGAAGAGACTCACGTAGGACGGTCA -3′; ASO-Vκ3-5’ss: 5′-CCAGTGGTATCTGAAATTGGAAATA-3′) and an irrelevant ASO (ASO-Ctrl: 5′- CCTCTTACCTCAGTTACAATTTATA-3′) were designed and purchased at Gene Tools, LLC (Philomath, USA). Experiments were performed at 0.5 million cells/mL in the presence of 3 µM ASO. Culture samples and supernatants were harvested at day 1 or 2 for subsequent flow cytometry analysis or RNA/protein extraction, as described below. Supernatants of cells were collected and stored at −20°C until used for enzyme-linked immunosorbent assays (ELISAs).

### RT-PCR and PCR

RNA was extracted using the TRIzol™ reagent (Invitrogen) procedure. Reverse transcription was carried out on 1 μg of DNase I (Invitrogen)-treated RNA using a High-Capacity cDNA Reverse Transcription kit (Applied Biosystems). Priming for reverse transcription was done with random hexamers. To analyze exon skipping, PCRs were performed on cDNA using the Taq Core Kit (MP Biomedicals) and appropriate primer pairs (Lpart1λ hs cons for: 5’-ATGGCCTGGDYYVYDCTVYTYCT-3’; Cλ hs rev: 5’-CTCCCGGGTAGAAGTCACT-3’; Lpart1 Vκ3 hs for: 5’-ATGGAAGCCCCAGCGCAGCTT-3’; Cκ hs rev : 5’-GCGGGAAGATGAAGACAGAT-3’; Lpart1 Vκ1-33 mur for: 5’-ACGGCCCAGCATGGACATGAGGGTCCCTGCTC-3’, Cκ mur rev : 5’-GCACCTCCAGATGTTAACTGC-3’; D = A or G or C, Y = C or T, V = A or C or G). To determine nucleotide sequence of normal and alternative transcripts, PCRs were performed on cDNA using the Phusion® High-Fidelity DNA Polymerase (New England BioLabs) and appropriate primers. After purification of the RT-PCR products using the NucleoSpin Gel and PCR clean-up kit (Macherey-Nagel) according to the manufacturer’s instructions, sequencing was performed using the BigDye™ Terminator v3.1 Cycle Sequencing Kit on a 3130 × l Genetic Analyzer ABI PRISM (Applied Biosystems). Quantification of PCR products was also performed using an Agilent 2100 Bioanalyzer (Agilent Technologies) according to the Agilent High Sensitivity DNA kit instructions.

### ELISA

Culture supernatants and sera were analyzed for the presence of Igκ by ELISA. ELISAs were performed in polycarbonate 96-multiwell plates, coated overnight at 4°C (100 μl per well) with 2 μg/ml Igκ antibodies (Goat Anti-Human kappa-UNLB, SouthernBiotech, ref 2060-01) in PBS. After three successive washing steps in PBS with 0.05% Tween® 20 (Sigma-Aldrich), a blocking step with 100 μl of 3% bovine serum albumin (BSA) (Euromedex) in PBS was performed for 30 min at 37°C. After three washing steps, 50 μl of sera/supernatant or standards Igκ/Igλ (Southern Biotechnologies, 400 ng/ml) were diluted into successive wells in 1% BSA/PBS buffer and incubated for 2 h at 37°C. After three washing steps, 100 μl per well of 1 μg/l alkaline phosphatase (AP)-conjugated goat anti-human antibodies (Goat Anti-Human kappa-AP, SouthernBiotech, ref 2060-04) was incubated in PBS with 0.05% Tween® 20 for 2 h at 37°C. After three washing steps, AP activity was determined by adding 100 μl of substrate for AP (SIGMAFAST™ p-nitrophenyl phosphate tablets, Sigma-Aldrich) during ∼15 min, the reaction was then blocked with the addition of 50 μl of 3 M NaOH (Sigma-Aldrich). Optic density was measured at 405 nm on a Multiskan FC microplate photometer (Thermo Scientific).

### Western blot

Cells were lysed in radioimmunoprecipitation assay (RIPA) Lysis and Extraction Buffer (Thermo Scientific) supplemented with protease and phosphatase inhibitor cocktail. Lysates were sonicated and protein levels were quantified by Pierce™ BCA Protein Assay Kit (Thermo Scientific). Proteins were denatured at 94°C for 5 min before separation on SDS-PAGE (4–20% Mini-PROTEAN® TGX™ Precast Protein Gels were used (Bio-Rad Laboratories)). Proteins were then electro-transferred onto Trans Blot Turbo polyvinylidene fluoride membranes (Bio-Rad Laboratories). Western blots were probed with Goat Anti-Human lambda-UNLB (SouthernBiotech, ref 2070-01) or Goat Anti-Human kappa-UNLB (SouthernBiotech, ref 2060-01) antibodies. Detection was performed using an HRP-linked rabbit anti-goat secondary antibody and chemiluminescence detection kit (ECL Plus™, GE Healthcare) using ChemiDoc™ Touch Imaging System (Bio-Rad Laboratories). Image Lab™ Software (Bio-Rad Laboratories) was used for relative quantification of the bands.

### Cell viability assays

Cell viability was assessed by flow cytometry after staining with 7-AAD (BD Pharmingen, ref 559925). Briefly, 500,000 cells were incubated for 15 min with 7-AAD (200 μl) in PBS supplemented with 2% FCS and 2 mM (ethylenedinitrilo) tetraacetic acid (EDTA) (Sigma-Aldrich), at 4 °C in the dark. Data were acquired on a BD Pharmingen Fortessa LSR2 (BD Biosciences, San Jose, CA, USA) and analyzed using Flowlogic™ software (Miltenyi Biotec).

### Engraftment of SK-MM-2 myeloma cells in Rag2^-/-^γC^-/-^ mice

3–4-mo-old mice were used in all experiments and maintained in our animal facilities, at 21–23°C with a 12-h light/dark cycle. Experiments were performed according to the guidelines of the ethics committee in Animal Experimentation of Limousin (registered by the National Committee). SK-MM-2 were harvested during the exponential growth phase (5 x 10^5^ cells/ml), mixed with Matrigel (Becton Dickinson) and engrafted by subcutaneous injection in previously shaved Rag2^-/-^γC^-/-^ mice (2 x 10^6^ cells/mice) . The mice were palpated every 2 days. When the tumour became detectable, two perpendicular diameters were measured and the tumor size was calculated using the elipsa volume formula. When the tumour reached ∼100mm^3^, mice received 50 µl intratumoral injections of ASO (ASO-Vκ3-5’ss or ASO-Ctrl: 25 nmoles/injection) or PBS, 4 times every 2-3 days. The size of the tumour was then measured daily and serum was taken every 4 days. Mice were ethically sacrificed when tumor volumes reached ∼800-900mm^3^ as represented in the death curve.

### LPS stimulation of mouse B cells and ASO treatments

Splenic B cells isolated from C57BL/6, mice were purified with the EasySep Mouse B Cell Isolation Kit (Stemcell Technologies). B cells were cultured for 4 days in RPMI 1640 with UltraGlutamine (Lonza) containing 10% fetal calf serum (FCS) (Dominique Dutscher), 1 mM sodium pyruvate (Eurobio), 1% AANE (Eurobio), 50 U/ml penicillin/50 μg/ml streptomycin (Gibco), and 129 μM 2-mercaptoethanol (Sigma-Aldrich). Splenic B cells were stimulated with either 1 μg/ml lipopolysaccharide (LPS) (LPS-EB Ultrapure, InvivoGen) for 4 days.

For ASO treatments, vivo-morpholino ASOs (ASO-Jκ1-5’ss ASO: 5′-ACTTTGGATTCTACTTACGTTTGAT-3′; ASO-JH2-5’ss ASO: 5′-AGAGGTTGTAAGGACTCACCTGA-3′) and an irrelevant ASO (ASO control: 5′-CCTCTTACCTCAGTTACAATTTATA-3′) were designed and purchased at Gene Tools, LLC (Philomath, USA). Stimulated splenic B cells were cultured in the presence of 3 μM ASO for 3 days, starting from day 1 of LPS stimulation. Culture samples were harvested at day 4 for subsequent RNA extraction and analysis by RT-PCR and RACE-PCR.

### RNA sequencing analysis

RNA-seq were performed on the Illumina NextSeq500 and analyzed with DESeq2 statistical analysis at Nice-Sophia-Antipolis Functional Genomics Platform. The resulting matrix of gene-level raw counts were used with DESeq2 (Love et al., 2014) to extract differentially expressed (DE) genes (adjusted p value < 0.05 and absolute log2foldchange ≥ 0.75; **Table S1**) for comparison and downstream analysis. Ingenuity Pathway Analysis (IPA) software was used to represent as heat map specific genes involved in cell death and different proteostasis pathways.

### High-throughput sequencing of immunoglobulin repertoire

The clonal abundance was estimated after high-throughput sequencing of VDJ junctions. Repertoire sequencing was performed as previously described (Li et al., 2013). Briefly, RNA (100-500 ng) was extracted from cells, and 5’RACE-PCR (rapid amplification of 5’ cDNA-ends by polymerase chain reaction) was performed using CH1γ or Cκ specific reverse primers (Boice et al., 2016). Sequencing adapter sequences were added by primer extension. The amplicons were sequenced on an Illumina MiSeq sequencing system using the MiSeq kit Reagent V3 600 cycles. Repertoire analysis was performed using the IMGT/HighV-QUEST tool and the R software (Alamyar et al., 2012; Bischof and Ibrahim, 2016). Briefly, CDR3 junctions were identified using HighV-QUEST. Based on these annotations, the reads were grouped into clonotypes that shared the same CDR3 and V, D (for heavy chains) and J sequences. To define the extent of variability allowed between CDR3, each CDR3 amino acid sequence was aligned against itself to define a maximum alignment score. Then, all CDR3 bearing the same V(D)J segments were aligned against each other. CDR3 were considered as coming from the same clonotype if their alignment score was superior or equal to 70% of their maximum alignment score. Then, the relative abundance of each clonotype was calculated. For this study, clones are defined either by a single read or by several identical reads, which makes it possible to consider all clones.

### Statistical Analysis

Results are expressed as means ± SEM, and overall differences between variables were evaluated by a Student’s t test using Prism GraphPad software (San Diego, CA)

## ACKNOWLEDGMENTS

The authors thank the staff of the Biologie Intégrative Santé Chimie Environnement (BISCEm) technical platforms at the University of Limoges (animal and cell cytometry facilities). This work was supported by grants from INCa (PLBIO2022-110 to LD), ANR (2017-CE15-0024-01 to LD), Ligue Contre le Cancer (comités Corrèze, Haute-Vienne), Fondation Française pour la Recherche contre le Myélome et les Gammapathies monoclonales (FFRMG) and Région Nouvelle-Aquitaine (AAP ESR 2020 and 2024 to LD). JML was funded by Ministry of Higher Education, Research and Innovation (MESRI) and Ligue Contre le Cancer fellowships.

