## Supplementary material for "Selective killing of plasma cell clones using splice-switching oligonucleotides targeting immunoglobulin variable exons": SUPLLEMENTAL TEXT

**SUPPLEMENTARY INFORMATION**

**Figure S1: Sequence of Igλ transcripts expressed in RPMI 8226 and XG6 myeloma cells**

**(A-B)** Representation of the Igλ sequence from RPMI 8226 **(A)** and XG6 **(B)** cells. Sequences were aligned using the IMGT-VQUEST software and cryptic donor splice sites involved in alternative splicing are shown in red, with predictive splicing scores noted below. ASO hybridization are also represented (blue), and in **(B)** the asterisk indicates the mismatch identified for Igλ sequence from XG6 cells.

**Figure S2: Decreased Igκ production after treatment of SK-MM-2 cells with ASO-Vκ3-3'ss.**

SK-MM-2 cells were untreated (-) or treated with ASO-Vκ_3-15_-3'ss (ASO) or an irrelevant control ASO (Ctrl) for 48 hours. **(A)** Igκ light chain protein levels were analyzed by Western Blot after normalization to the expression of β-actin. **(B)** ELISA assays were performed on culture supernatants to determine total Igκ amounts. Data from n=5-6 experiments are expressed as mean ± SEM and unpaired two-tailed Student's t test was used to determine statistical significance. (*P < 0.05,**P < 0.01, ***P < 0.001, ****P < 0.0001).

**TABLE S1: List of differentially expressed genes after treatment of SK-MM-2 cells with ASO-V**κ**_3-15_-3'ss or control ASO.** (abs(log2FC) ≥ 0,75); pvalue Adj < 0,05).
