## Supplementary figures and images for "Selective killing of plasma cell clones using splice-switching oligonucleotides targeting immunoglobulin variable exons"

### SUPPLEMENTAL FIGURES

#### Slide 1
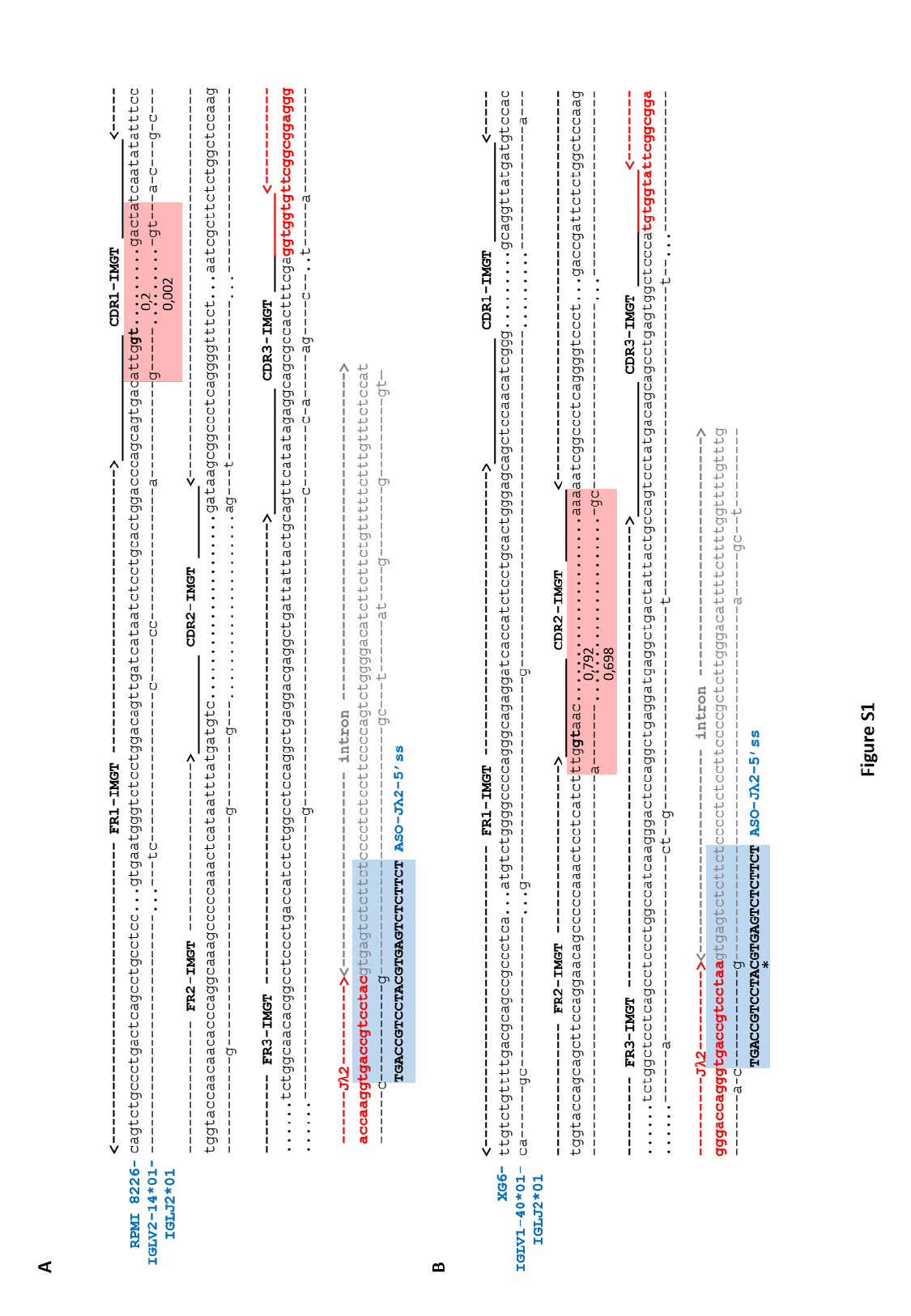

#### Slide 2
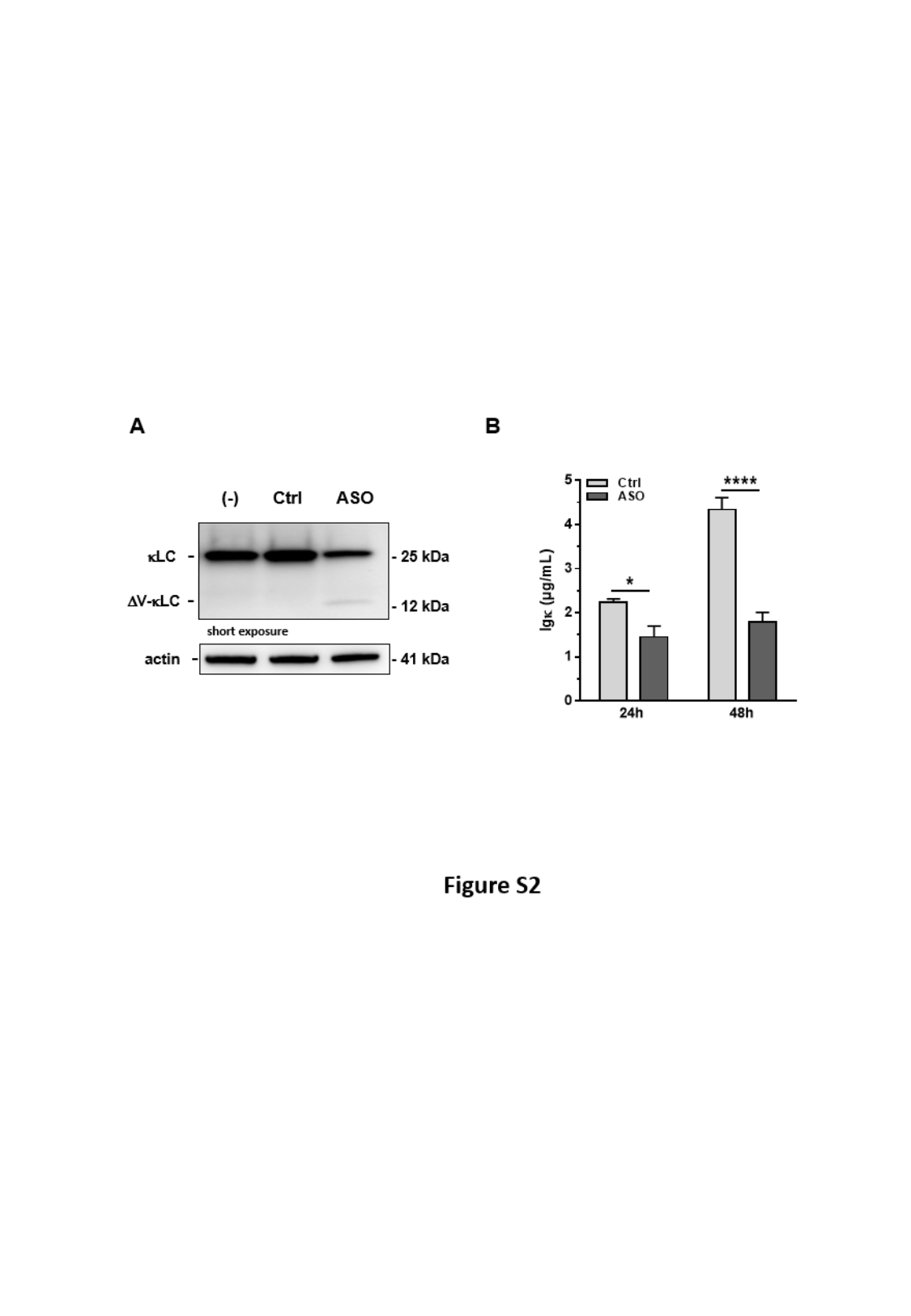
